# Resident myeloid-derived immune cells contribute to early lipopolysaccharide-induced cytokine secretion in mouse soleus muscle

**DOI:** 10.64898/2026.06.23.734036

**Authors:** Finleigh P. Fitton, Deborah A. Morse, Kevin J. Cusack, Bryce J. Gambino, Thomas L. Clanton

## Abstract

Skeletal muscles secrete a variety of cytokines in response to inflammatory stimuli such as lipopolysaccharide (LPS); however, the contributions of resident macrophages or other non-muscle cells to the secretory responses are not well understood. Here we tested the LPS responsiveness of mouse soleus when a critical toll receptor adapter protein (Myd88) was knocked down only in myeloid-derived cells (e.g. resident macrophages). The phenotype is referred to as LyzMyd88^−/−^ ; the litter mate controls were Myd88^fl/fl^. In solei from LyzMyd88^−/−^mice, cytokine secretory rates for interleukin-6 (IL-6) and keratinocyte-derived cytokine (KC, CXCL1) were significantly reduced to 56.3%, and 60.6% of control, respectively, over the first hour of LPS exposure. In the second hour, the secretion of granulocyte colony stimulating factor (G-CSF), IL-6, KC(CXCL1) and monocyte chemoattractant protein-1 (MCP-1, CCL2) were greatly elevated by 5-10-fold in both phenotypes compared to the first hour. However, only MCP-1 secretion was decreased to 70.6% of control in the second hour. We also tested the secretory response to buffer containing 1% sterile mouse plasma because dilute plasma is known to amplify the responses of macrophages to LPS. Treatment with 1% plasma alone affected baseline measures of some cytokines but resulted in no further increases in secretion during either hour of exposure. However, small and gradual increases in secretory rates were observed for several cytokines over the study period, with or without plasma, with the largest responses seen in IL-6 and KC. Overall, the results are consistent with a significant early contribution of myeloid-derived, resident immune cells to the cytokine secretory responses of intact oxidative skeletal muscle. In addition, small quantities of plasma in the buffer have no independent stimulatory effects on cytokine secretion.

## Introduction

The ability of both in vitro and in vivo skeletal muscle to secrete cytokines in response to inflammatory stimuli has been well established for over 20 years [1–3]. Extensive evidence has also documented the ability of isolated skeletal muscle fibers or myogenic cells to secrete immune cytokines [1,2,4]. However, there are many unanswered questions as to the relative contribution of muscle fibers vs. other muscle phenotypes that are resident within intact muscle.

In this report we first test the hypothesis that addition of low levels of sterile mouse plasma provides an additional and independent stimulus for inflammatory cytokine production from skeletal muscle. In a previous publication we demonstrated that 1% plasma in buffer vs. buffer alone amplified cytokine secretory responsiveness to LPS in isolated mouse soleus and in extensor digitorum longus muscle [5]. The results paralleled previous reports of the effects of low levels of plasma on cytokine secretion in isolated macrophages from humans [6]. However, whether there were independent (i.e. without LPS) influences of plasma on secretory rates or if there were time-dependent effects of the isolation procedures and exposure to oxygenated buffer were not tested.

In another series of experiments, we tested the hypothesis that a significant component of the inflammatory cytokine secretory responses in isolated whole muscle arises from resident myeloid-derived immune cells, such as tissue macrophages. To test this hypothesis, we utilized a well-established myeloid cell-specific knockdown mouse that targets “myeloid differentiation primary response protein 88” (Myd88), found only in myeloid derived cells [7–9]. Myeloid-derived cells include macrophages, monocytes, dendritic cells, neutrophils, eosinophils, basophils and mast cells. Resident macrophages are of the highest concentration among these immune cells in muscle and therefore they were the primary target for Myd88 knockdown [10]. Myd88 serves as a critical adapter protein for all toll-like receptors (TLRs) that reside on the plasma membrane that are sensitive to inflammatory mediators such as lipopolysaccharides (LPS), as well as other damage associated- or pathogen associated-molecular patterns [11–13]. It is also a critical adapter for interleukin-1 beta (IL1-β) receptor activation, an important pro-inflammatory cytokine [14]. Animal models of global Myd88 knockout exhibit almost no plasma cytokine responsiveness to LPS injections [11]. Whole animal myeloid cell-specific knockout of Myd88 reduces circulating inflammatory cytokines to less than half of the values of wildtype controls [7]. Therefore, this transgenic model provides a powerful tool for estimating the contribution of resident myeloid cells to inflammatory signaling in isolated tissues.

Our previous studies in intact mice have demonstrated marked sex differences in the whole animal inflammatory responses to sepsis when specific inflammatory pathways are knocked down, only in skeletal muscle [15,16]. Whether these differences reflect responses at the skeletal muscle level or at some other level of organization of the immune system is not known. Therefore, we also sought to evaluate how sex affects cytokine secretion at both the whole skeletal muscle fiber level and at the level of resident myeloid inflammatory cells.

## Methods

### Animal Care and Handing

This study was carried out in strict accordance with the recommendations in the Guide for the Care and Use of Laboratory Animals of the National Institutes of Health. All protocols were approved by the University of Florida Institutional Animal Use Committee (IACUC #202200185). All procedures met the requirements of ARRIVE guidelines 2.0 [17]. C57BL/6 mice were ordered from The Jackson Laboratory (strain #000664) at 12-15 weeks of age and group housed until euthanasia and muscle collection. The transgenic colony was bred, in-house, as described below. Mice were housed within University of Florida Animal Care Facilities in a 12:12-h light-dark cycle at 20-22°C / 30-60% relative humidity and maintained on Teklad 2918 rodent diet (Envigo, Indianapolis, IN). All animals were transported to the laboratory from a University of Florida vivarium on the morning of muscle collection.

### Myeloid-specific Myd88 knock down

For knockout experiments, mutant mice, Myd88 floxed mice (strain #008888) were bred with LysMCre mice (strain #004781). Both strains were purchased from The Jackson Laboratory (Bar Harbor ME). LysMCre mice have Cre-recombinase expression controlled by the promoter on the lysozyme M gene, which is expressed only in myeloid-derived cells (macrophages, basophils, etc). The mice were then cross-bred to produce male and female offspring with the LyzCre-Myd88^flox/flox^ genotype (Fig. 1a). The myeloid cells of these offspring have Myd88 continuously knocked down throughout life. Results were compared to a control group of litter mates carrying only the Myd88^flox/flox^ transgene, without the LyzCre transgene. This is not a new model as it has been used previously for similar purposes by multiple studies [7–9] Every mouse was individually genotyped using established primers from Jax labs for both Myd88 and Lyz-Cre. The use of the primers has been previously demonstrated along with validation of the effects of knockdown on mRNA and protein expression in isolated tissue macrophages [7–9].

**Figure 1.**
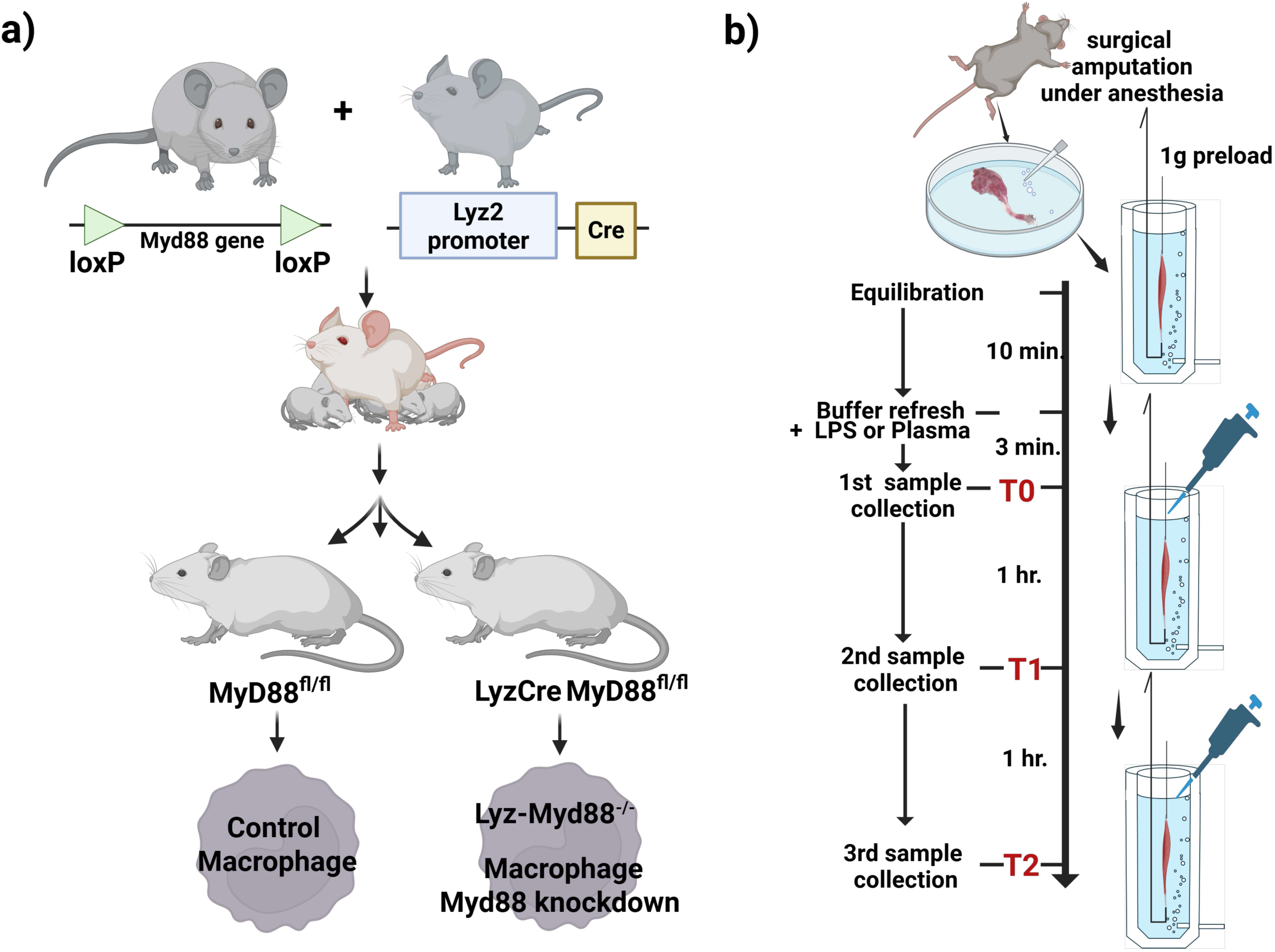
**a)** Breeding of myeloid cell-specific Myd88 knockout mice. Mice with loxP site flanking the Myd88 gene were bred to mice with Cre-recombinase inserted into the lysozyme 2 gene promoter (Lyz2). The floxed Myd88^+/+^ offspring mice were used as litter-mate controls. Lyz2 is the particular Lyz isoform uniquely expressed in myeloid cells. Cre recombinase is then only expressed in macrophages and other myeloid-derived immune cells, causing Myd88 to be knocked down only in myeloid cells during or after hematopoiesis. Myd88 is a critical adapter protein for most toll-like receptors (TLR) and for interleukin-1 receptor. **b)** The experimental protocol design as described in text. Created in Biorender, Inc.

### Experimental Design

The mice were anesthetized under isoflurane anesthesia (0.6 L/min O_2_, 4% isoflurane for induction, 1.5% for maintenance). Then both hind limbs were rapidly removed and placed under oxygenated, preheated Krebs buffer (Fig. 1b). Mice were immediately euthanized by cervical dislocation and separation of the heart from the great vessels. Soleus muscles were chosen for these experiments because of their faster accessibility for isolation, thus reducing the mechanical stress on the muscle, and because of the fact that they have a larger number of inflammatory macrophages per muscle volume compared to fast muscles [18,19]. In addition, in previous work we demonstrated very similar cytokine secretory rates per dry mass in isolated extensor digitorum longus (EDL) and soleus muscles [5]. Hind limbs were skinned to keep the muscles as clean as possible and free from hair throughout isolation (Figure 1b). Both muscles from each animal were individually isolated by a pair of experienced surgeons to ensure equal isolation times. Krebs Ringer buffer solution, equilibrated with 95% O_2_/5% CO_2_, was used for all isolation procedures and for bath incubation: (in mmol/L) 0.45 Na_2_SO_4_, 0.6 Na_2_HPO_4_, 1.0 MgCl_2_, 5.9 KCl, 2.0 CaCl_2_, 21.0 NaHCO_3_, 121.0 NaCl, and 11.5 glucose. The buffer was filtered with sterile 0.2 μm PES Nalgene filters (ThermoFisher, Waltham, MA #597-4520) to restrict bacteria, endotoxin and other particles. Excised solei were secured with suture on both tendons and hung in 3 mL oxygenated tissue baths (Radnoti, Covina, Calif, #158303) containing 2.5 mL of buffer at 35°C. The muscles were maintained with a preload of 1 g throughout the study, a value resulting in a resting length that was just below optimum length. Solei were never electrically stimulated during the protocols. Once in the baths, the solei were acclimated to the oxygenated buffer for 10 min. Then the buffer was removed and replaced with fresh buffer as described below.

To study the effects of supplemental 1% plasma, adult male and female wild type C57BL/6 mice were utilized. After equilibration, the solei were exposed to 2.5 ml of fresh Krebs buffer (3 ml baths). In one half of the muscles, 25 μL of commercial mouse plasma (Innovative research sIGMS-C57-N, Novi MI) was added and in the other half of muscles (controls) an additional 25 μL of fresh Krebs buffer was added. Prior to use, the mouse plasma was treated with an endotoxin removal kit (BioRad, Hercules, CA #PUR030), aliquoted and frozen at −20°C for single use for each experiment. Bath samples of 200 μL were removed at 3 time points, 3 min after the first fresh buffer replacement (T0), 1 hour post (T1), and 2 hours post (T2) plasma exposure (Fig. 1b). To each collected sample, an aliquot of a protease inhibitor cocktail was added (Bimake, Houston, TX #B14001) prior to flash freezing in liquid nitrogen and storing at - 80C° C for later analyses. Following the experiment, the solei were removed from the muscle baths, dried overnight on a slide warmer at 37°C and weighed for normalizing cytokine secretion to dry muscle weight.

To study the effects of LPS in the muscles in response to knockdown of myeloid Myd88, the experiments were conducted identically to the plasma exposure study, except that all muscles were exposed to 1% plasma throughout the study (Valley Biomedical #AP3054, Winchester, VA) and 1 μg/mL LPS (Sigma-Aldrich, St. Louis, Mo #L4515) was added to both baths at the time of the first buffer wash (Fig. 1b). The solei in the baths were randomly selected from either the LyzCre-Myd88^−/−^ mice or their litter mate Myd88^flox/flox^ controls. The experimental group assignment was not blinded during the experiment but was blinded during analysis. An additional group of eight muscles, four with plasma and four without, were tested for endotoxin contamination at the T2 sample collection time using a Limulus Amebocyte Lysate assay, (LAL assay kit, ToxinSensor™ Chromogenic LAL Endotoxin Assay Kit – GenScript, Piscataway, NJ). The primary outcome variables were always the amount of cytokine produced at a give timepoint in the experiment or within a specific time interval.

### Multiplex cytokine analysis

Cytokine analyses from frozen bath samples were performed using a Luminex MAGPIX multiplex system (Luminex Corporation, Austin TX) employing a Millipore MAP Mouse Cytokine/Chemokine Magnetic Bead Panel #MCYTOMAG-70K (Millipore-Sigma, Burlington, MA). Of the available cytokines for that panel, only a subset of 16 analytes were chosen based on the results of our previous studies using this preparation, with and without 1% plasma or LPS [5,20]. These cytokines included granulocyte colony-stimulating factor (G-CSF), interferon gamma (INF-γ), interleukin 1α (IL-1α), interleukin 1β (IL-1β), interleukin 6 (IL-6), interleukin 10 (IL-10), interleukin 13 (IL-13), interleukin 15 (IL-15), interferon gamma-induced protein (IP-10), keratinocytes-derived chemokine (KC/CXCL1), monocyte chemoattractant protein (MCP-1/CCL2), macrophage inflammatory protein-1 α (MIP1-α), macrophage inflammatory protein-1 β (MIP1-β), macrophage inflammatory protein-2 (MIP-2), regulated upon activation normal T cell expressed and secreted (RANTES), and tumor necrosis factor-α (TNFα). Analytes in which the average concentrations were below the minimum limit of detection (LOD) at any point in the collection period were assumed to be equal to the LOD for that analyte. Only analytes that displayed a significant average cytokine concentration above the LOD for either set of experiments at any time point were reported in the results section.

### Statistical Analysis

Power analyses to determine a minimum sample size was performed using G*Power 3.1 [21]. IL-6 measurement outcomes were used as the primary outcome variable. For the LyzMyd88^−/−^ experiments, we sought a power of 0.8, an α < 0.05, a minimum acceptable difference of 25%, resulting in an overall effect size of 1.21, which estimated a minimum sample size of 7. For the effects of plasma, we expected smaller responses with less variance and used a minimum meaningful difference of 40% with an effect size of 2.2 resulting in a minimum acceptable sample size of 5. No outcome data were removed or excluded from any groups reported except because of surgical isolation errors or technical difficulties during preparation or analyses, such as failure of the Luminex detection assay.

All statistics were generated using SAS JMP 18.0.2. For each cytokine studied for its response to 1% plasma, a mixed Model, 3 -way ANCOVA with repeated measures was used. The covariate factors of interest were plasma treatment (Tx), time (interval 1 or 2), and sex, with the crossed effects for Time * Tx. The repeated measures factor was handled as individual muscle name, entered as a random nominal variable. Preplanned comparisons of the effects of plasma at each time point, for each cytokine, were evaluated as pairwise contrasts. A nearly identical statistical model was used for LyzMyd88 knockdown experiments, except the treatment variable was “phenotype.” In many analyses, the capacity to detect differences was improved and the datasets made more parametric by transforming the cytokine measures into natural logarithms (ln). Some changes in [cytokine] within an interval were negative numbers. To apply Ln transformations to negative numbers, a fixed integer, larger than the most negative value within a group was added to all of the sample values to be compared, based on standard approaches [22]. The ANCOVA for both studies demonstrated that there were no differences in secretion rates due to SEX. Therefore, for final graphical displays, we combined the data for both sexes and reported the *P* value for the effects of SEX as a separate covariate outcome, while graphically displaying the responses of the two sexes with different symbols. Because of the high variance within many of the data sets, we graphically reported the central tendency and variance as medians ± 25-75% quartiles.

## Results

### Effects of 1% plasma on soleus cytokine secretion

This experiment was used to test the hypothesis that in isolated muscle, exposure to 1% mouse plasma (in the absence of other inflammatory stimuli) has a direct stimulatory effect on cytokine secretion. An example of the concentration data collected for this experiment is displayed in Fig. 2 for IL-6 in the tissue baths over the two-hour exposure in either buffer control (Fig. 2a and 2c) or 1% plasma in buffer (Fig 2b or 2d). The IL-6 concentrations are expressed per mg dry muscle mass. There were differences in concentrations in the bath, during interval one interval two in all groups, but no statistically differences in between responses with and without plasma in each interval or between males and females.

**Figure 2.**
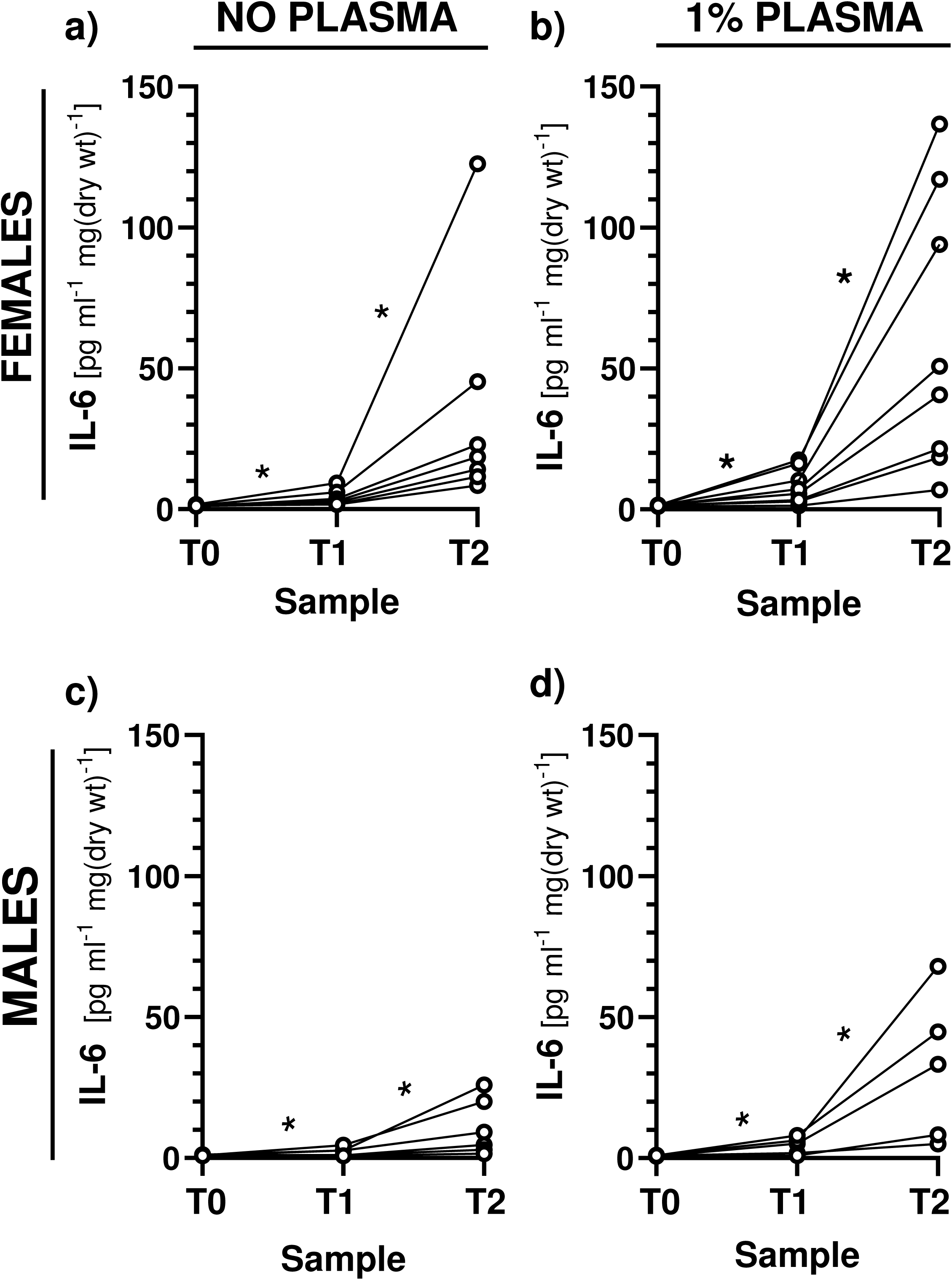
Representative data from the experiments showing IL-6 concentrations (mg/ml) in the baths expressed as concentration per mg dry soleus. Effects of plasma supplementation in the tissue baths at each time point (T0-T2). a) response in female solei, without plasma (*n* = 7), b) response of female solei with 1% plasma (*n =* 8), c) male response without plasma (*n* = 6), d) male response without plasma (*n* = 5). * *P* <0.05, paired *t* test. Each value indicates data from single muscle from single mouse. Time T0 = 3 minutes post plasma supplementation. Time T1 = 1 hour post, time T2 = two hours post. Mean differences between groups were tested between sexes; sex had no significant effects on changes in concentration between sample times.

We based our primary statistical findings on the total amount of each cytokine secreted within each time interval (i.e. during interval 1: T1-T0 and during interval 2: T2-T1), effectively removing baseline effects on these measurements (Fig. 3). Grouped secretion rates were corrected for muscle dry weight and dilution effects in the bath. Only the cytokines, among all those studied (listed in methods), that showed significant responsiveness above their LOD are included in the analyses. Because no effect of sex could be identified for any cytokines using univariate comparisons, the data for both sexes were combined and the influence of the sex (nominal variable) handled as a covariate in ANCOVA. The *P* values for SEX and other independent covariates in the ANOVA are listed in Table 1.

**Figure 3.**
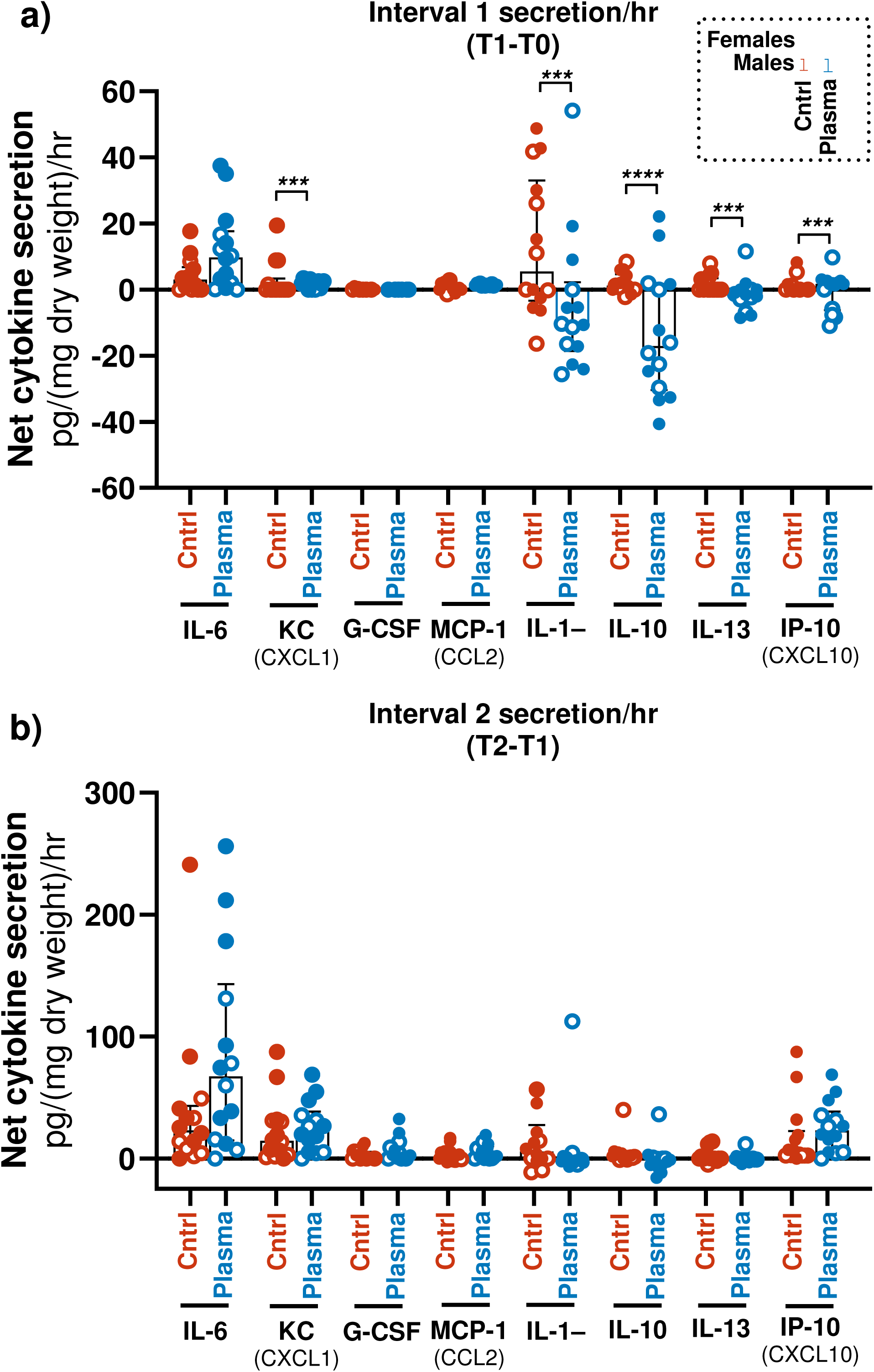
**a)** Whole soleus cytokine secretion within the first hour: Interval 1 (T1-T0). **b)** results in the second hour, Interval 2 (T2-T1). All data are normalized to bath volume and dry soleus mass. Multiway, repeated measures ANCOVA results are tabulated in Table 1. Central tendency was expressed as median ± 25-75% quartiles. Male and female data are designated in closed or open symbols, respectively, and combined within groups because there was no univariate influence of sex. Each value indicates data from a single muscle and from a single mouse. No plasma control: *n* = 14; plasma effects: *n* = 14. * = *P* represents preplanned pairwise contrasts of the effects of plasma vs. no plasma, post ANCOVA. **** *P=*<0.0001, ***= *P*<0.001, **= *P*<0.01, * *P*<0.05. Effects of time on the secretion between interval 1 and 2 in panel 3b for all responsive cytokines: ΘΘΘΘ = <0.0001, ΘΘΘ=<0.001, ΘΘ<0.01, Θ =<0.05, independent of plasma.

**Table 1.** Results of multiway ANCOVA for effects of 1% plasma and covariates.

| Cytokine | P value<br>Whole model | P value<br>Plasma Treat. | P value<br>Time | P value<br>Treat. x Time<br>(Time interval) | P value<br>Repeated<br>Meas. | P value<br>Sex | Dependent Var.<br>Transformation |
| --- | --- | --- | --- | --- | --- | --- | --- |
| IL-6 | <0.0001 | 0.101 | <0.0001 | 0.815 | 0.001 | 0.484 | Ln |
| KC | <0.0001 | 0.062 | <0.0001 | 0.004 | 0.002 | 0.146 | Ln |
| GCSF (CXCL1) | 0.011 | 0.136 | 0.001 | 0.100 | 0.001 | 0.924 | None |
| MCP-1 (CCL2) | 0.045 | 0.308 | 0.007 | 0.451 | 0.955 | 0.380 | None |
| IL-1α | 0.010 | 0.002 | 0.002 | 0.012 | 0.112 | 0.779 | Ln |
| IL-10 | 0.001 | 0.004 | 0.019 | 0.036 | 0.974 | 0.276 | Ln |
| IL-13 | 0.047 | 0.023 | 0.161 | 0.134 | 0.638 | 0.448 | Ln |
| IP-10 (CXCL10) | 0.007 | 0.047 | 0.008 | 0.089 | 0.463 | 0.239 | Ln |
| IL-15 | 0.070 | 0.624 | 0.031 | 0.053 | 0.156 | 0.694 | None |
| MIP-1α (CCL3) | 0.522 | 0.400 | 0.785 | 0.218 | 0.544 | 0.338 | Ln |

Model: Predicted [Cytokine] = $\mu$ + Plasma Tx + Time + Sex +(Plasm Tx. \* Time) + Repeated measures Time (interval 1 vs. interval 2).
Dependent variable transformation: Ln = natural log transformation prior to ANCOVA
Post hoc analyses expressed on Figure 3.
All Benjamini-Hochberg corrected $P < 0.12$ for cytokines in which the Whole Model was $P < 0.05$

Our original hypothesis was that plasma exposure alone would increase inflammatory cytokine production over time. As shown in Fig. 3a, only KC (CXCL1) showed a very small but statistically significant increase in secretion within interval 1. Over interval 2, Fig. 3b, no cytokines exhibited any increase in secretion rate in response plasma treatment compared to no plasma controls. Interestingly, four cytokines (IL-1α, IL-10, IL-13 and IP-10) showed significantly reduced ‘apparent’ secretion in interval 1 (Fig. 3a) but not interval 2 (Fig. 3b). “Apparent” refers here to the probability that some cytokine at time T0 (Supplementary Table S1), may be affected by processes other than secretion, such as loss to plasma soluble receptors, degradation, binding to the tissue or glassware or other factors. All cytokines exhibited in Fig. 3, except IL-13, showed an overall increase in cytokine secretion over time (i.e. Θ Symbols Fig. 3b), regardless of whether they received 1% plasma treatment. This suggests that isolation and exposure of the solei to the oxygenated bath environment for sufficient periods, affects secretion. Effects of 1% plasma on absolute concentration/ dry weight at each specific time point (T0,T1,T2) is shown in the supplemental Table S1, which emphasizes the small but significant influences of plasma on some baseline cytokines.

To determine if some of the effects of plasma could have arisen from endotoxin contamination from muscle isolation procedures or contact to associated equipment, buffer samples were taken after solei were incubated in for the full 2 hrs, with or without 1% plasma (*n=* 4 per group). The results averaged 0.36 ± 0.16 SD, and 0.41 ± 0.08 EU/ml of endotoxin (∼0.5 ng/ml), with or without plasma, respectively (*P*=0.58). Based on previous dose-response curves for this same endotoxin formulation and isolated muscle protocol, these concentrations would not have impacted net cytokine secretion from mouse soleus over the collection period [5].

### Effects of LPS and Lyz-Myd88^−/−^ on soleus cytokine secretion

An example of the effects of LPS exposure on IL-6 bath concentrations over time (normalized to dry weight) in Myd88 control vs. LyzMyD88^−/−^ muscles is exhibited in Fig. 4. Note: the scale differences between Fig. 2 and Fig. 4, showing the overall large stimulatory effect of LPS on secretion rates. In all intervals and groups, LPS treatment stimulated secretion. There were no differences between secretory rates of IL-6 between females and males (not shown) when handled as univariate paired measurements.

**Figure 4.**
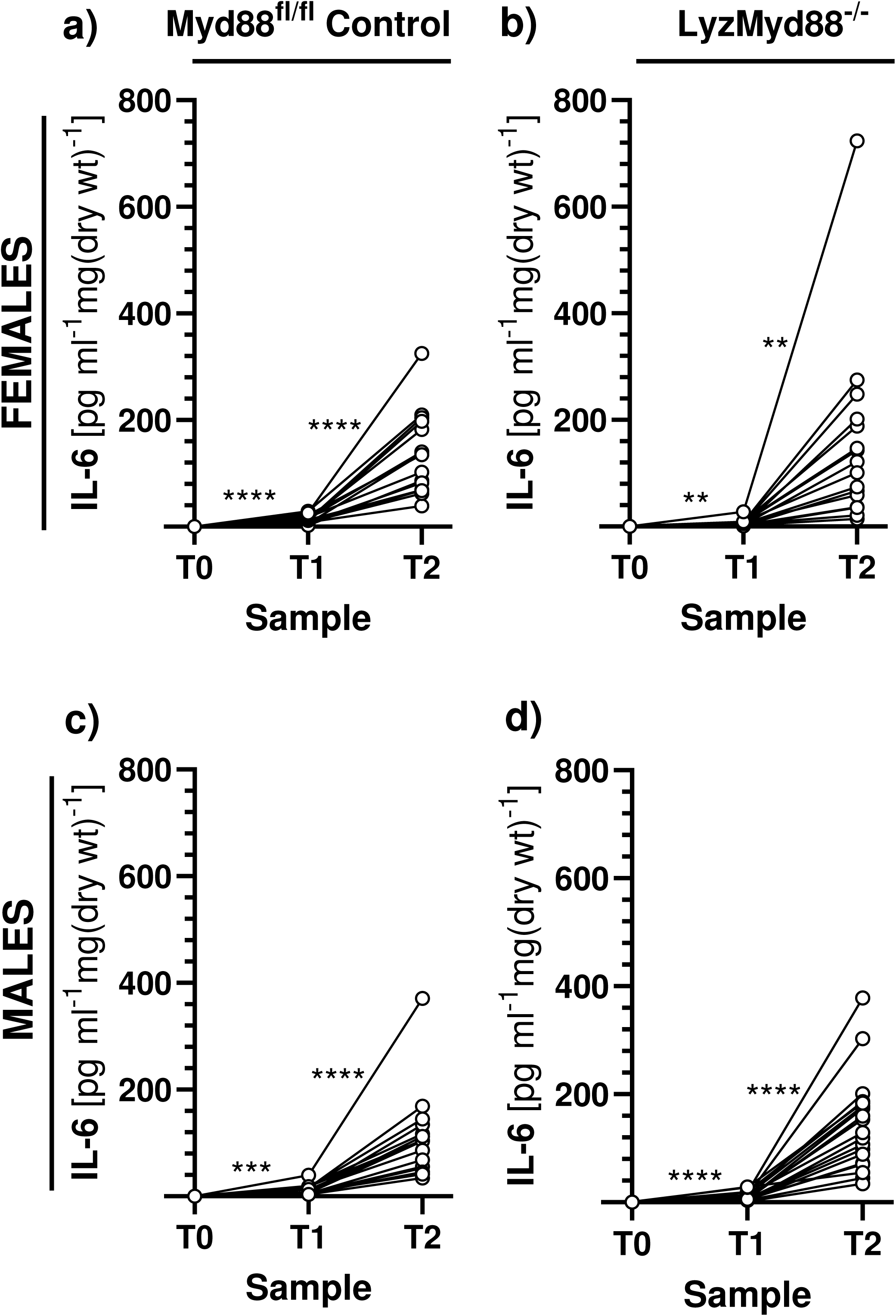
Representative data for IL-6 concentrations per mg dry soleus measured in the tissue baths in response to lipopolysaccharide (LPS), with and without myeloid Myd88 knockdown. **a)** response in female solei in Myd88^fl/fl^ control mice (*n* = 16), **b)** response of female solei in Lyz-Myd88^−/−^ muscles : (*n =19*), **c)** male response in control Myd88^fl/fl^ (*n* =16) **d)** male response in Lyz-Myd88^−/−^ mice (*n =19*). Each value indicates data from single muscle from single mouse. Sample T0 = 3 minutes post LPS supplementation. Sample T1 = 1 hour post; sample T2 = 2 hours post. ** = *P*<0.01, ***= *P*<0.001, ****= *P*<0.0001, paired t test. There were no significant differences between male and female responses in each interval, within genotype (using non-paired T test of changes within each interval).

Fig. 5 (a and b) exhibits the rates of secretion of all responsive cytokines to LPS within intervals one and two. Note, LPS did not induce a minimum detectable concentration for many cytokines within any time point studied. As described in Methods, these cytokines were not included in the final analysis or graphics. As with the effects of plasma, there were no significant effects of the covariate, SEX, on cytokine secretion (Table 2) and therefore, for the final analysis and graphical representation, the data from both sexes were combined into single groups with sex represented within each group as different symbols. The only cytokines that responded overall to LPS within intervals 1 and 2 were IL-6, KC(CXCL1), GCSF, and MCP-1(CCL2) (Fig. 5a and b). See Table 2 for *P* values for covariates of ANCOVA for this experiment. Within interval 1, the effects of myeloid-specific knockdown of Myd88 significantly reduced secretion of IL-6 in response to LPS to an average of 56.4% of control and reduced secretion of KC(CXCL1) to an average of 60.6% of control (Fig. 5a). There were no significant effects of knockdown in interval 2, except for MCP-1(CCL2), which reduced secretion, on average, to 70.6% of control. Time effects for overall secretion rate in response to LPS between interval 1 and 2 are represented by symbols (Θ) in Fig. 5b; LPS accelerated secretion rate of all responding cytokines in the second compared to the first hour.

**Figure 5.**
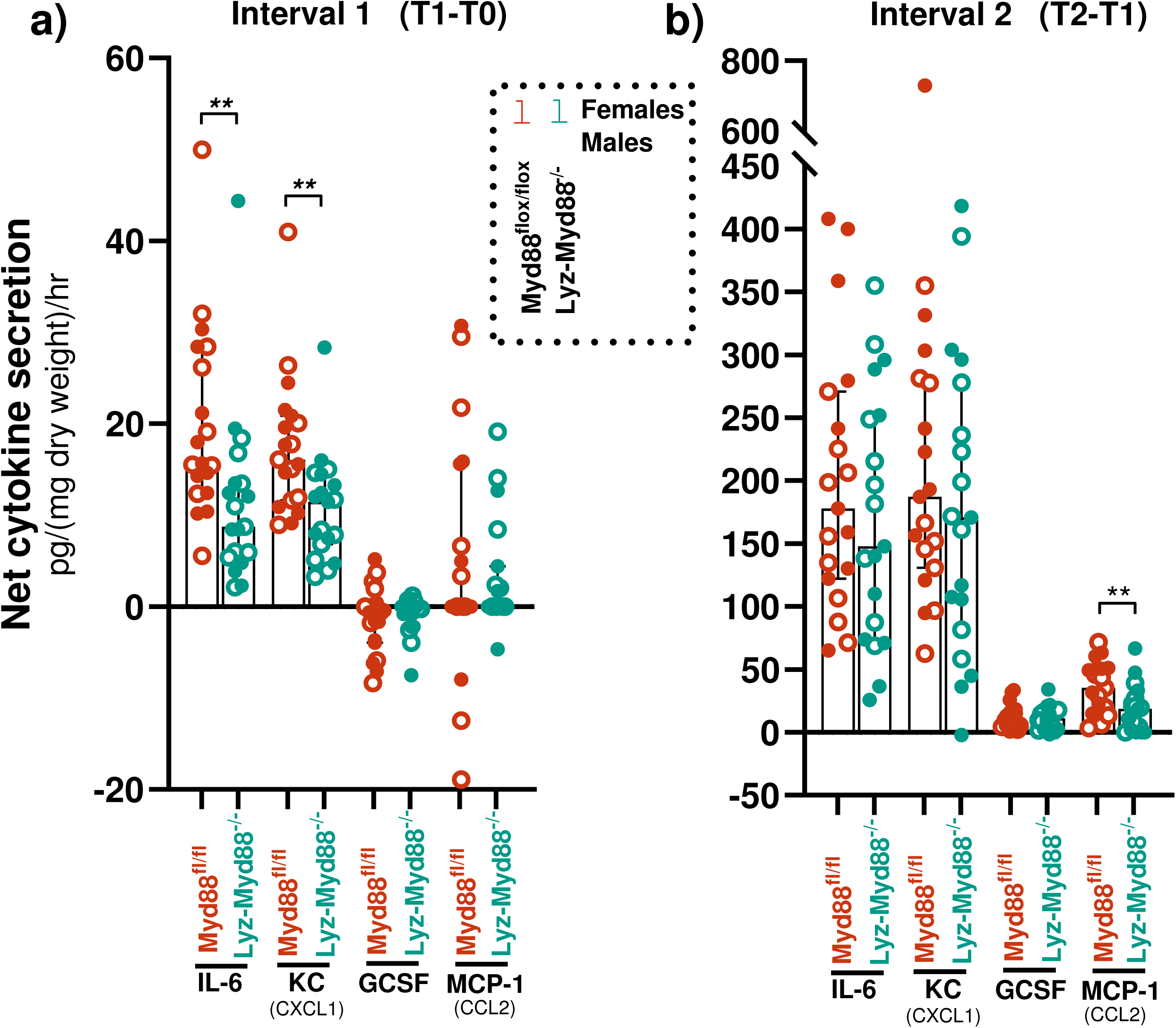
**a)** Cytokine secretion rates in response to LPS within interval 1 (T_1_-T_0_). **b)** cytokine secretion within interval 2 (T_2_-T_1_) from isolated solei in 1% protein buffer. All data normalized to bath volume and dry soleus weight. Total *n* in each group = 19. Central tendency expressed as median ± 25-75% quartiles. Males and females were combined, because there were no significant univariate effects of sex (See Table 2 for ANCOVA results for sex). Each value indicates data from a single muscle from a single mouse. Effects of phenotype (LyzMyd88 knockout) **= *P*<0.01, * = *P*<0.05. Effects of time, ANCOVA variable, on secretion rate between interval 1 and 2 in panel 5b: ΘΘΘΘ = <0.0001, independent of phenotype.

**Table 2.**
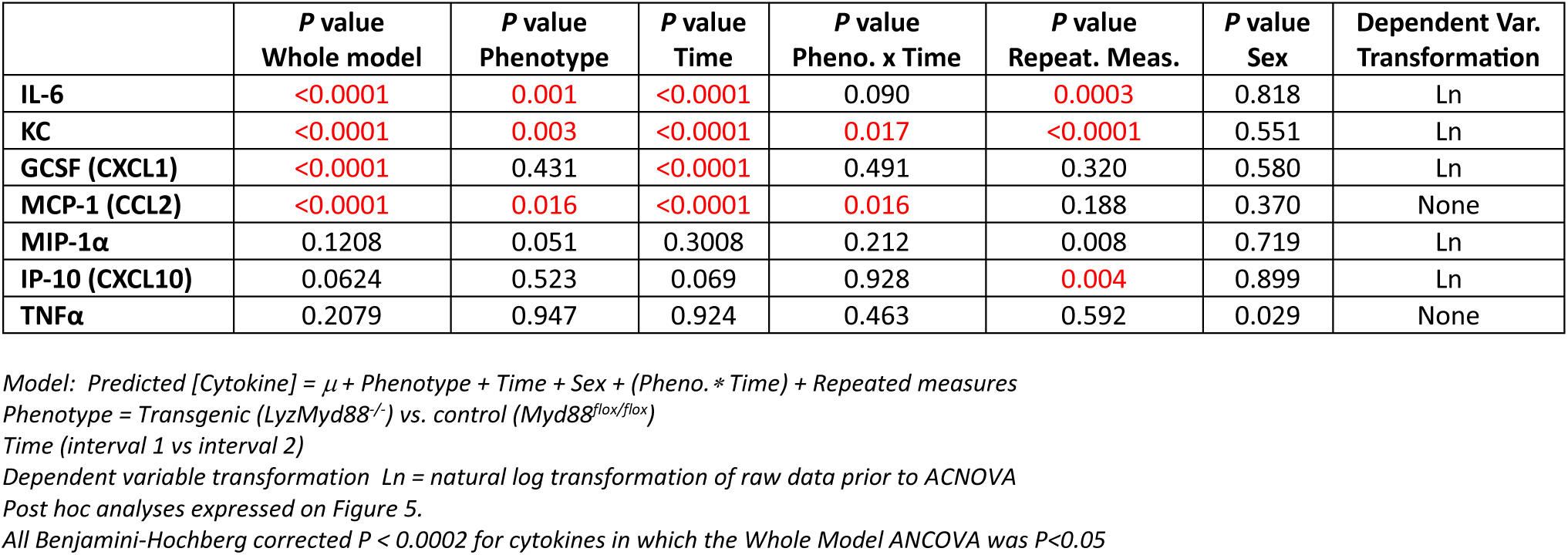
Results of multiway ANCOVA for LPS-stimulated muscle and covariates.

|  | *P* value<br>Whole model | *P* value<br>Phenotype | *P* value<br>Time | *P* value<br>Pheno. x Time | *P* value<br>Repeat. Meas. | *P* value<br>Sex | Dependent Var.<br>Transformation |
| --- | --- | --- | --- | --- | --- | --- | --- |
| IL-6 | <0.0001 | 0.001 | <0.0001 | 0.090 | 0.0003 | 0.818 | Ln |
| KC | <0.0001 | 0.003 | <0.0001 | 0.017 | <0.0001 | 0.551 | Ln |
| GCSF (CXCL1) | <0.0001 | 0.431 | <0.0001 | 0.491 | 0.320 | 0.580 | Ln |
| MCP-1 (CCL2) | <0.0001 | 0.016 | <0.0001 | 0.016 | 0.188 | 0.370 | None |
| MIP-1α | 0.1208 | 0.051 | 0.3008 | 0.212 | 0.008 | 0.719 | Ln |
| IP-10 (CXCL10) | 0.0624 | 0.523 | 0.069 | 0.928 | 0.004 | 0.899 | Ln |
| TNFα | 0.2079 | 0.947 | 0.924 | 0.463 | 0.592 | 0.029 | None |

Model: Predicted [Cytokine] = $\mu$ + Phenotype + Time + Sex + (Pheno. \* Time) + Repeated measures
Phenotype = Transgenic (LyzMyd88<sup>-/-</sup>) vs. control (Myd88<sup>flax/flax</sup>)
Time (interval 1 vs interval 2)
Dependent variable transformation Ln = natural log transformation of raw data prior to ANCOVA
Post hoc analyses expressed on Figure 5.
All Benjamini-Hochberg corrected $P < 0.0002$ for cytokines in which the Whole Model ANCOVA was $P < 0.05$

To further explore the time dependence of cytokine measurements in the bath, the absolute cytokine concentrations per dry soleus weight are expressed and compared at each collection time point, T0, T1 and T2 (Supplemental Table S2).

## Discussion

Our results are consistent with the following observations: resident tissue myeloid-derived cells contribute a significant proportion of the ‘early’ cytokine secretory responses to LPS (i.e. over the first hour), for IL-6 and KC/ CXCL1, and with a significant reduction in MCP-1 in the 2^nd^ hour. In separate experiments, 1% sterile mouse plasma had no direct stimulatory effect on cytokine secretion in the absence of LPS. However, for some cytokines, the addition of plasma either reduced or increased absolute cytokine values, possibly due to residual cytokines in the plasma supplement, cytokine adherence to soluble receptors in the plasma supplement [5], cytokine protein binding or degradation. The results also show that tissue isolation alone and/or exposure to the oxygenated buffer environment is an independent stimulus for significant cytokine secretion over a two-hour experimental time window.

### Contributions of myeloid cells

We speculate that the early contribution of myeloid-derived cells to overall cytokine secretion of the intact muscle is much higher than the value we report here. First, based on previous studies using exactly the same genetic model, Myd88 RNA and protein were only reduced in mature macrophages by ∼75% and 67%, respectively in the knockdown animals [7]. Secondly, once inflammatory cells, myofibers or other resident cells begin making significant concentrations of other inflammatory cytokines, these can provide a delayed, positive, secondary feedback to stimulate secretion via Myd88-independent mechanisms, e.g. TNFα or MCP-1. There are also alternative inflammatory pathways linked to toll receptors that can be stimulated via LPS that are not Myd88-dependent, most notably the “Toll/interleukin-1 receptor domain-containing adapter-inducing interferon-β” (TRIF) pathway [23]. However, recent work has demonstrated that TRIF knockdown does not suppress NFκB signaling in response to LPS in mouse macrophages when measured within the first 24 hour period of exposure [23,24]. It does cause a small inhibition of the inflammatory response at 48 hours of exposure to LPS [24]. Therefore, we concluded that this alternative pathway of TLR-induced inflammatory signaling is not relevant to the time-course of this specific study design.

Skeletal muscle is comprised of many cell phenotypes besides muscle fibers, including resident macrophages, muscle satellite cells, fibroblasts, adipocytes, regulatory T cells and endothelium [25–27]. Many of these are capable of producing immune cytokines in response to LPS, particularly myofibroblasts [1], adipocytes [28] and resident macrophages [29]. Specific resident myeloid-derived immune cells that could have been affected by Lyz-Myd88^−/−^ include macrophages, basophils, mast cells and dendritic cells. Estimates of resident macrophage populations in mouse muscle vary widely depending on the method and tissue studied. They are often reported as density/square area or in units common to flow cytometry samples. Both approaches are difficult to accurately interpret in terms total macrophage density per volume of intact muscle. However, some studies, using immunohistochemical imaging, have shown that macrophages comprise ∼10% of the total nuclear content of muscle in young mouse vastus intermedius (a slow soleus-like muscle) [10]. In rat solei, macrophages are found at a density of ∼6,900 cells/mm^3^ [18], or about 50,000 macrophages per soleus if macrophage density in rats can be applied to mice. Of these, only about 30% are designated as having the classic “M1-phenotype”, i.e. the polarization state considered pro-inflammatory and responsible for the inflammatory cytokine release in response to LPS [30]. Many muscle macrophages reside in the perimysium and epimysium of muscle [31], which would have a more rapid access to dissolved LPS in the tissue bath and could accelerate the overall secretory response of the intact muscle in the first hour of exposure, the time interval that appears most sensitive to manipulation of Lyz-Myd88 in our results.

### Effects of 1% plasma on cytokine secretion

Based on previous work in isolated muscle, adding 1% plasma to the tissue baths amplifies the skeletal muscle responsiveness to LPS [5], much like what happens in isolated human macrophages [6]. We hypothesized, a priori, that low concentrations of plasma might also have an independent effect in the absence of LPS. Our reasoning was based on a speculation that large molecular weight plasma proteins (e.g. soluble CD5, immunoglobulin or oxidized albumin [35]) may be recognized within the muscle microenvironment as ‘foreign’, resulting in an independent inflammatory stimulus. However, we found no apparent stimulation by plasma exposure alone on secretion of any cytokine species.

### Other Effects

Interestingly, exposure to the oxygenated tissue bath, with or without 1% plasma, significantly elevated the concentrations of two major muscle cytokines, IL-6 and KC/CXCL1 over the 2-hour exposure. Though it is unlikely the muscles experienced significant passive tension in the baths, it is possible that they may have been inadvertently stretched during isolation. Passive stretch is a unique stimulus for cytokine secretion [36]. Another possibility is the stress of 95% O_2_ exposure or exposure to the buffer environment could have constituted a stimulus. In our previous work in isolated rat diaphragm, we observed a reduction in muscle total glutathione by ∼50% over the course of a one hour of exposure to a oxygenated tissue bath [37], suggesting the tissue bath environment itself is an ongoing oxidative stress. Oxidative stress can stimulate IL-6 secretion, in part by activating protein degradation pathways [38,39], and IL-6 is thought to be able to independently induce secretion of KC(CXCL1) in stress conditions [40,41].

### Potential effects of the immune network in the muscle microenvironment

There are many potential inflammatory signaling interactions between different phenotypes within the muscle microenvironment. For example, a secondary response of macrophages to inflammatory signaling is local shedding of soluble receptors for inflammatory cytokines such as soluble TNF receptor (sTNFR), soluble IL-1 receptors (sIL-1R), and soluble IL-6 receptor (sIL-6 receptor) [42–44]. This occurs in the presence of inflammatory mediators such as LPS. These are also expected to be elevated in the plasma supplement. Interestingly, TNFα and IL-1β were noticeably absent or extremely low in measurements taken from the tissue baths during LPS exposure. Since they are considered extremely important in normal inflammatory responses to LPS in all tissues, their absence in the tissue bath was puzzling. It is possible, however, that TNFα was still an active participant within the microenvironment of the muscle but constrained to operate in the ‘bound state’ through binding to soluble TNF receptors and subsequent formation of receptor clustering on plasma membranes [45]. This would make TNFα unavailable for diffusion out of the isolated muscle interstitial space and into the bath. Another possibility for why we saw little IL-β or TNFα secretion into the bath is that IL-6 (presumably in very high concentration within the interstitium) has a paradoxical inhibitory effect on pro-inflammatory cytokine expression [46,47]. These possibilities suggest that the immune network found within skeletal muscle is highly interconnected and responds to stressors in a complex and dynamic manner, particularly within the highly condensed and localized intercellular compartment.

### Experimental limitations

One important limitation to this work is a lack of biological validation of the LyzMyd88 knockdown model in myeloid cells. As mentioned, every mouse utilized was validated by genotyping for both Myd88^fl/fl^ and Lyz-Cre genes using published PCR oligonucleotides as primers [7–9]. In addition, the effectiveness of identical knockout model used in other studies on gene transcription and protein expression were previously well documented in macrophages or monocytes from different tissues [7–9]. Nevertheless, the reader should be cautioned that there is always a possibility of genetic drift in the germ lines that could affect the outcomes.

A second consideration has to do with the interpretation of the data. Although the major results are expressed in terms of “secretion rates” (Figs. 3 and 5) within one-hour intervals, secretion rates could be much higher and faster than actually measured in the highly diluted tissue baths. Proteins such as cytokines, when secreted from individual cells have a circuitous diffusion path within the interstitium and out into the surrounding tissue bath resulting in a slow accumulation of secretory products. Furthermore, the total interstitial volume of the mouse soleus in young mice is estimated to be ∼20 μL [48] compared to a tissue bath volume of 2,500 μL. If the intact muscle were secreting a specific cytokine at a constant rate, e.g. 10 pg per minute per soleus, it would require 375 minutes to accumulate 10 pg/ml in the bath, not accounting for diffusion delays within the interstitium. Based on ‘effective’ diffusion constant estimates for molecular weight molecules in the category of cytokines [49], it would require many minutes for the cytokines to actually exit the muscle, depending on the location of the secreting cells (note: the soleus is roughly 0.5 mm in radius). Therefore, caution should be used in terms of interpretation of the timing of the responses in the intact, non-perfused muscle, which inherently distorts the actual kinetics of production at a cellular level.

Other myeloid cells could also have contributed significantly to cytokine secretion. However, dendritic cells residing in normal skeletal muscle are largely plasmacytoid, anti-inflammatory, positioned for viral surveillance and unlikely to be impacted by LPS [32]. Basophils are only substantially recruited into muscle during inflammation, with very few resident cells in non-inflamed muscle [33]. Mast cells are resident, particularly in blood vessels, and capable of secreting inflammatory cytokines but are rare, being only ∼1 cell per mm^2^ in normal C57BL/6 mouse soleus [34]. Therefore, we propose that it is unlikely that other myeloid phenotypes contribute substantially to whole muscle cytokine secretory response in these isolated muscle experiments.

### Implications and ideas for future work

In two recent reports, our laboratory evaluated the effects of conditional knockout of specific inflammatory pathways within muscle fibers, i.e. muscle fiber-specific Myd88 knockout [15] and muscle fiber-specific IL-6 knockout [16]. In both studies, immune responsiveness of the whole organism was greatly suppressed during septic shock. The results of this study provide some additional insights into understanding of how muscle can influence whole-body immune regulation. It suggests that any changes in muscle fiber cytokine secretion could interact with resident inflammatory cells or vice versa. One interaction that could be longer lasting than the two-hour window of this experimental design is the capacity for macrophages to change polarization states. The primary tissue macrophage phenotype in muscle is anti-inflammatory, i.e. various subtypes of M2 polarization [50]. However, M2 macrophages can transform to a pro-inflammatory, M1, phenotypes over time in response to inflammatory signals like LPS [50] making their potential contributions in the inflammatory responses of the whole organism likely to be much larger when evaluated over extended periods. Since muscle makes up 40% of body mass, these contributions alone could have marked impacts on circulating cytokines and overall immune regulation. More work is needed to determine if these or other interactive mechanisms within the myoimmune axis are inherent components of innate or acquired immunity. This area of research remains an investigative space that may lead to new therapeutic strategies for recovery from infection, sepsis or injury [51,52].

## Acknowledgements

**Grants:** This work was supported by NIH R01GM118895-0 (TLC) and the BK and Betty Stevens Endowment (TLC).

The Figure 1 was created using Biorender.com under publication license.

## Current addresses of authors when different from title page

Bryce Gambino: Liberty University Medical School, 1971 University Boulevard, Lynchburg, VA 24515

Finleigh P. Fitton: University of North Florida, Physical Therapy Ph.D. Program, Brooks College of Health, Jacksonville, FL, 32224

## Data availability Statement

Add data used for this publication will be archived in a publicly available site once the manuscript is accepted for publication.

## Author contributions

Finleigh P. Fitton: performed experiments and data analysis, wrote 1^st^ draft of the manuscript and approved final manuscript.

Deborah A. Morse: performed experiments, a major role in data acquisition and analysis, contributed to the manuscript and approved the final draft.

Kevin J. Cusack: assisted in design of the experiments, performed experiments, performed data analysis and approved the final manuscript.

Bryce J. Gambino: performed experiments and approved the final manuscript.

Thomas L. Clanton: designed experiments, performed data analysis, contributed to final draft of manuscript.

## Supplementary Material

Tables S1 and S2 can be found at https://figshare.com/articles/dataset/Supplementary_Tables_S1_and_S2/28635407?file=53135549

## Supplementary Data and Text

**Table S1.**
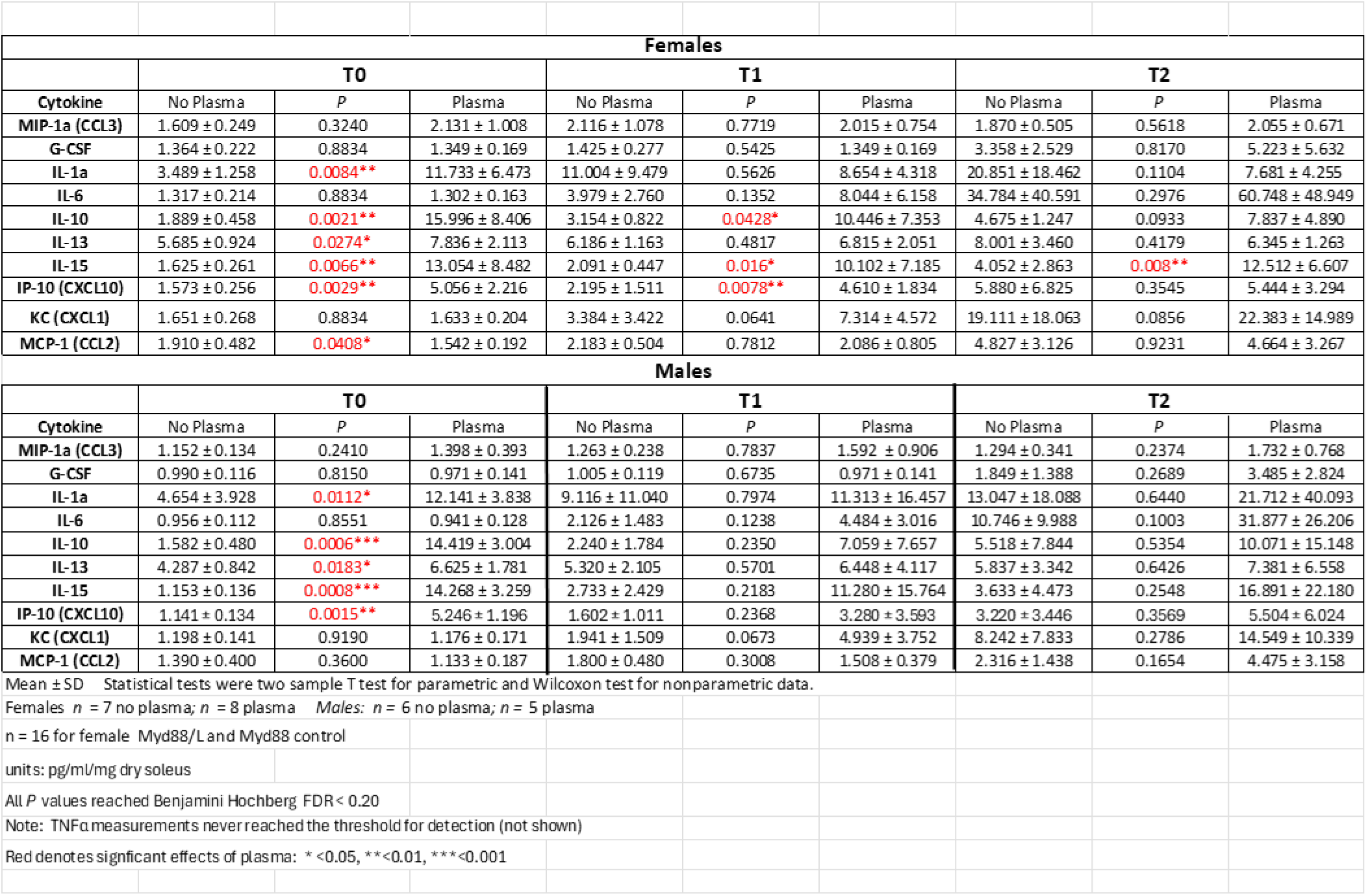
Concentrations of cytokines in the muscle baths at the three points of sample collection in the 1% plasma experiments. A number of cytokine measurements exhibited elevated baseline concentrations in both males and females in the presence of plasma compared to no plasma (Table S1; time, T_0_). In both males and females, IL1-α, IL-10, IL-13, IL-15 and IP10 were elevated. Since these samples were taken ∼3 min after refreshing the buffer/plasma mixture in the bath, they are not likely to represent secretion, but rather their presence in the 1% commercial plasma supplement or on the muscle surfaces.

**Table S2.**
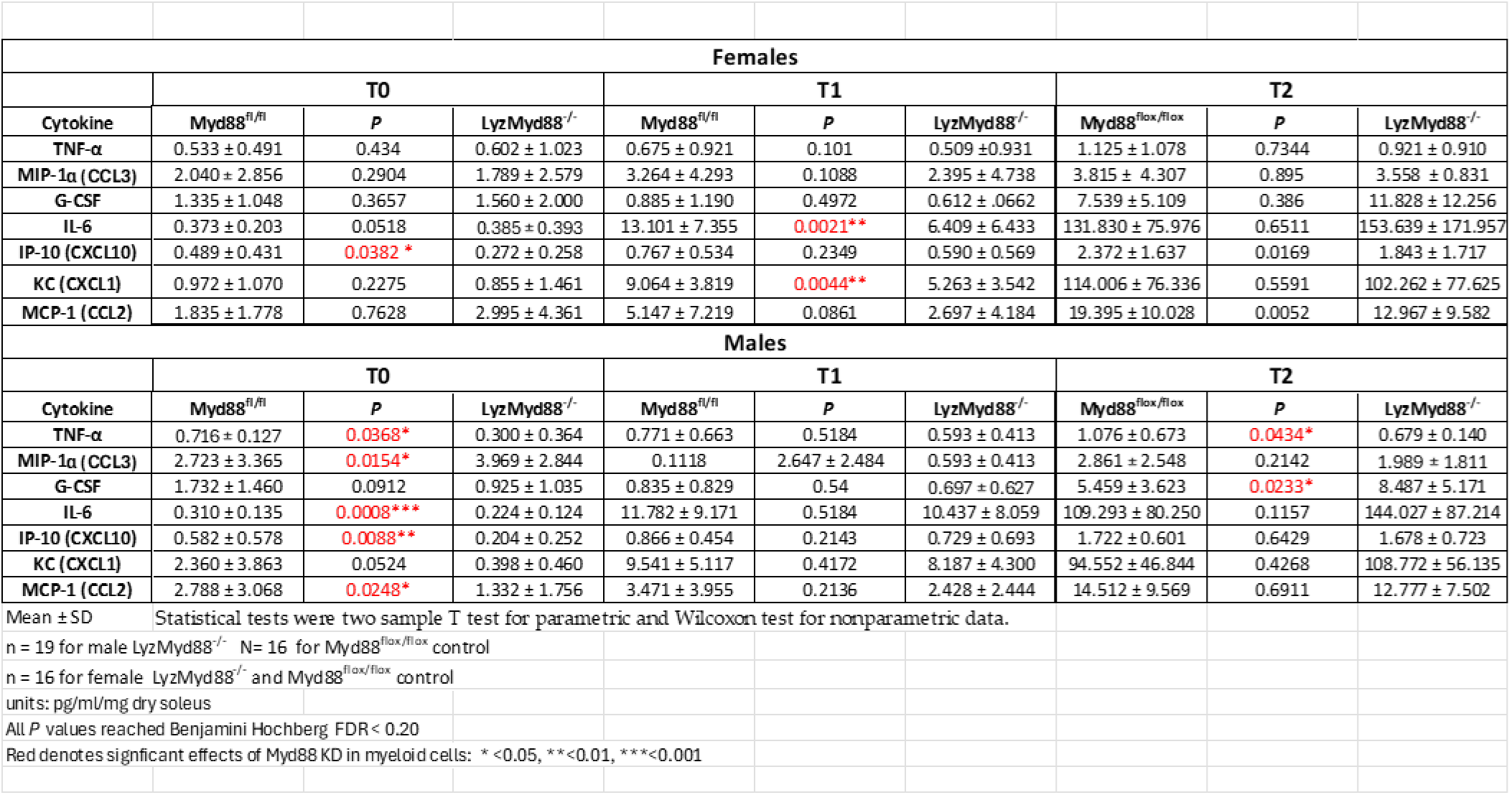
Concentrations of cytokines in the muscle baths at the three points of sample collection in the LPS stimulation experiments. In general, in this experiment, baseline values of cytokines at T0 were extremely low. When comparing the absolute concentrations of muscle cytokines within the baths at each time point, females showed a reduction in IP-10 and KC secretion in the LyzMyd88 KO muscles at T_2_ and decreases in IL-6 and KC at T_1_. They also showed reductions in MCP-1 and IP-10 at T_2_. In contrast, males exhibited reductions in TNFα, IL-6, IP10, and MCP1 at T_0._ At T_2_, TNFα was marginally decreased and G-CSF marginally increased in the LYZ-Myd88^−/−^ muscles. Muscles from LyzMyd88^−/−^mice produced lower baseline cytokines (particularly in males) in the KO mice suggesting that some of the baseline cytokine measurements observed may have arisen from the muscle tissue during the brief < 3 min of buffer exposure, and that these were influenced by LyzMyd88^−/−^. None of these single timepoint measurements affected the data reported in the main text which were based on the rate of secretion within intervals.

