## Supplementary material for "Resident myeloid-derived immune cells contribute to early lipopolysaccharide-induced cytokine secretion in mouse soleus muscle": S1

### Supplementary Data and Text

**Table S1. Effects of plasma exposure vs buffer alone on measured cytokines at each sample collection time**

| Females | | | | | | | | | |
|---|---|---|---|---|---|---|---|---|---|
| | T0 | | | T1 | | | T2 | | |
| Cytokine | No Plasma | *P* | Plasma | No Plasma | *P* | Plasma | No Plasma | *P* | Plasma |
| MIP-1a (CCL3) | 1.609 ± 0.249 | 0.3240 | 2.131 ± 1.008 | 2.116 ± 1.078 | 0.7719 | 2.015 ± 0.754 | 1.870 ± 0.505 | 0.5618 | 2.055 ± 0.671 |
| G-CSF | 1.364 ± 0.222 | 0.8834 | 1.349 ± 0.169 | 1.425 ± 0.277 | 0.5425 | 1.349 ± 0.169 | 3.358 ± 2.529 | 0.8170 | 5.223 ± 5.632 |
| IL-1a | 3.489 ± 1.258 | 0.0084** | 11.733 ± 6.473 | 11.004 ± 9.479 | 0.5626 | 8.654 ± 4.318 | 20.851 ± 18.462 | 0.1104 | 7.681 ± 4.255 |
| IL-6 | 1.317 ± 0.214 | 0.8834 | 1.302 ± 0.163 | 3.979 ± 2.760 | 0.1352 | 8.044 ± 6.158 | 34.784 ± 40.591 | 0.2976 | 60.748 ± 48.949 |
| IL-10 | 1.889 ± 0.458 | 0.0021** | 15.996 ± 8.406 | 3.154 ± 0.822 | 0.0428* | 10.446 ± 7.353 | 4.675 ± 1.247 | 0.0933 | 7.837 ± 4.890 |
| IL-13 | 5.685 ± 0.924 | 0.0274* | 7.836 ± 2.113 | 6.186 ± 1.163 | 0.4817 | 6.815 ± 2.051 | 8.001 ± 3.460 | 0.4179 | 6.345 ± 1.263 |
| IL-15 | 1.625 ± 0.261 | 0.0066** | 13.054 ± 8.482 | 2.091 ± 0.447 | 0.016* | 10.102 ± 7.185 | 4.052 ± 2.863 | 0.008** | 12.512 ± 6.607 |
| IP-10 (CXCL10) | 1.573 ± 0.256 | 0.0029** | 5.056 ± 2.216 | 2.195 ± 1.511 | 0.0078** | 4.610 ± 1.834 | 5.880 ± 6.825 | 0.3545 | 5.444 ± 3.294 |
| KC (CXCL1) | 1.651 ± 0.268 | 0.8834 | 1.633 ± 0.204 | 3.384 ± 3.422 | 0.0641 | 7.314 ± 4.572 | 19.111 ± 18.063 | 0.0856 | 22.383 ± 14.989 |
| MCP-1 (CCL2) | 1.910 ± 0.482 | 0.0408* | 1.542 ± 0.192 | 2.183 ± 0.504 | 0.7812 | 2.086 ± 0.805 | 4.827 ± 3.126 | 0.9231 | 4.664 ± 3.267 |
| **Males** | | | | | | | | | |
| | T0 | | | T1 | | | T2 | | |
| Cytokine | No Plasma | *P* | Plasma | No Plasma | *P* | Plasma | No Plasma | *P* | Plasma |
| MIP-1a (CCL3) | 1.152 ± 0.134 | 0.2410 | 1.398 ± 0.393 | 1.263 ± 0.238 | 0.7837 | 1.592 ± 0.906 | 1.294 ± 0.341 | 0.2374 | 1.732 ± 0.768 |
| G-CSF | 0.990 ± 0.116 | 0.8150 | 0.971 ± 0.141 | 1.005 ± 0.119 | 0.6735 | 0.971 ± 0.141 | 1.849 ± 1.388 | 0.2689 | 3.485 ± 2.824 |
| IL-1a | 4.654 ± 3.928 | 0.0112* | 12.141 ± 3.838 | 9.116 ± 11.040 | 0.7974 | 11.313 ± 16.457 | 13.047 ± 18.088 | 0.6440 | 21.712 ± 40.093 |
| IL-6 | 0.956 ± 0.112 | 0.8551 | 0.941 ± 0.128 | 2.126 ± 1.483 | 0.1238 | 4.484 ± 3.016 | 10.746 ± 9.988 | 0.1003 | 31.877 ± 26.206 |
| IL-10 | 1.582 ± 0.480 | 0.0006*** | 14.419 ± 3.004 | 2.240 ± 1.784 | 0.2350 | 7.059 ± 7.657 | 5.518 ± 7.844 | 0.5354 | 10.071 ± 15.148 |
| IL-13 | 4.287 ± 0.842 | 0.0183* | 6.625 ± 1.781 | 5.320 ± 2.105 | 0.5701 | 6.448 ± 4.117 | 5.837 ± 3.342 | 0.6426 | 7.381 ± 6.558 |
| IL-15 | 1.153 ± 0.136 | 0.0008*** | 14.268 ± 3.259 | 2.733 ± 2.429 | 0.2183 | 11.280 ± 15.764 | 3.633 ± 4.473 | 0.2548 | 16.891 ± 22.180 |
| IP-10 (CXCL10) | 1.141 ± 0.134 | 0.0015** | 5.246 ± 1.196 | 1.602 ± 1.011 | 0.2368 | 3.280 ± 3.593 | 3.220 ± 3.446 | 0.3569 | 5.504 ± 6.024 |
| KC (CXCL1) | 1.198 ± 0.141 | 0.9190 | 1.176 ± 0.171 | 1.941 ± 1.509 | 0.0673 | 4.939 ± 3.752 | 8.242 ± 7.833 | 0.2786 | 14.549 ± 10.339 |
| MCP-1 (CCL2) | 1.390 ± 0.400 | 0.3600 | 1.133 ± 0.187 | 1.800 ± 0.480 | 0.3008 | 1.508 ± 0.379 | 2.316 ± 1.438 | 0.1654 | 4.475 ± 3.158 |

Mean ± SD Statistical tests were two sample T test for parametric and Wilcoxon test for nonparametric data.

Females *n* = 7 no plasma; *n* = 8 plasma *Males: n* = 6 no plasma; *n* = 5 plasma

n = 16 for female Myd88/L and Myd88 control

units: pg/ml/mg dry soleus

All *P* values reached Benjamini Hochberg FDR < 0.20

Note: TNFα measurements never reached the threshold for detection (not shown)

Red denotes signficant effects of plasma: * <0.05, **<0.01, ***<0.001

**Table S1. Concentrations of cytokines in the muscle baths at the three points of sample collection in the 1% plasma experiments.** A number of cytokine measurements exhibited elevated baseline concentrations in both males and females in the presence of plasma compared to no plasma (Table S1; time, $T_0$). In both males and females, IL1-$\alpha$, IL-10, IL-13, IL-15 and IP10 were elevated. Since these samples were taken ~3 min after refreshing the buffer/plasma mixture in the bath, they are not likely to represent secretion, but rather their presence in the 1% commercial plasma supplement or on the muscle surfaces.

**Table S2. LPS-induced cytokines accumulated at each sample collection time in control vs. myeloid MYD88 Knockdown.**

| | | | | | | | | | |
|---|---|---|---|---|---|---|---|---|---|
| **Females** | | | | | | | | | |
| | **T0** | | | **T1** | | | **T2** | | |
| **Cytokine** | **Myd88$^{fl/fl}$** | ***P*** | **LyzMyd88$^{-/-}$** | **Myd88$^{fl/fl}$** | ***P*** | **LyzMyd88$^{-/-}$** | **Myd88$^{flox/flox}$** | ***P*** | **LyzMyd88$^{-/-}$** |
| **TNF-α** | 0.533 ± 0.491 | 0.434 | 0.602 ± 1.023 | 0.675 ± 0.921 | 0.101 | 0.509 ±0.931 | 1.125 ± 1.078 | 0.7344 | 0.921 ± 0.910 |
| **MIP-1α (CCL3)** | 2.040 ± 2.856 | 0.2904 | 1.789 ± 2.579 | 3.264 ± 4.293 | 0.1088 | 2.395 ± 4.738 | 3.815 ± 4.307 | 0.895 | 3.558 ± 0.831 |
| **G-CSF** | 1.335 ± 1.048 | 0.3657 | 1.560 ± 2.000 | 0.885 ± 1.190 | 0.4972 | 0.612 ± .0662 | 7.539 ± 5.109 | 0.386 | 11.828 ± 12.256 |
| **IL-6** | 0.373 ± 0.203 | 0.0518 | 0.385 ± 0.393 | 13.101 ± 7.355 | 0.0021** | 6.409 ± 6.433 | 131.830 ± 75.976 | 0.6511 | 153.639 ± 171.957 |
| **IP-10 (CXCL10)** | 0.489 ± 0.431 | 0.0382 * | 0.272 ± 0.258 | 0.767 ± 0.534 | 0.2349 | 0.590 ± 0.569 | 2.372 ± 1.637 | 0.0169 | 1.843 ± 1.717 |
| **KC (CXCL1)** | 0.972 ± 1.070 | 0.2275 | 0.855 ± 1.461 | 9.064 ± 3.819 | 0.0044** | 5.263 ± 3.542 | 114.006 ± 76.336 | 0.5591 | 102.262 ± 77.625 |
| **MCP-1 (CCL2)** | 1.835 ± 1.778 | 0.7628 | 2.995 ± 4.361 | 5.147 ± 7.219 | 0.0861 | 2.697 ± 4.184 | 19.395 ± 10.028 | 0.0052 | 12.967 ± 9.582 |
| **Males** | | | | | | | | | |
| | **T0** | | | **T1** | | | **T2** | | |
| **Cytokine** | **Myd88$^{fl/fl}$** | ***P*** | **LyzMyd88$^{-/-}$** | **Myd88$^{fl/fl}$** | ***P*** | **LyzMyd88$^{-/-}$** | **Myd88$^{flox/flox}$** | ***P*** | **LyzMyd88$^{-/-}$** |
| **TNF-α** | 0.716 ± 0.127 | 0.0368* | 0.300 ± 0.364 | 0.771 ± 0.663 | 0.5184 | 0.593 ± 0.413 | 1.076 ± 0.673 | 0.0434* | 0.679 ± 0.140 |
| **MIP-1α (CCL3)** | 2.723 ± 3.365 | 0.0154* | 3.969 ± 2.844 | 0.1118 | 2.647 ± 2.484 | 0.593 ± 0.413 | 2.861 ± 2.548 | 0.2142 | 1.989 ± 1.811 |
| **G-CSF** | 1.732 ± 1.460 | 0.0912 | 0.925 ± 1.035 | 0.835 ± 0.829 | 0.54 | 0.697 ± 0.627 | 5.459 ± 3.623 | 0.0233* | 8.487 ± 5.171 |
| **IL-6** | 0.310 ± 0.135 | 0.0008*** | 0.224 ± 0.124 | 11.782 ± 9.171 | 0.5184 | 10.437 ± 8.059 | 109.293 ± 80.250 | 0.1157 | 144.027 ± 87.214 |
| **IP-10 (CXCL10)** | 0.582 ± 0.578 | 0.0088** | 0.204 ± 0.252 | 0.866 ± 0.454 | 0.2143 | 0.729 ± 0.693 | 1.722 ± 0.601 | 0.6429 | 1.678 ± 0.723 |
| **KC (CXCL1)** | 2.360 ± 3.863 | 0.0524 | 0.398 ± 0.460 | 9.541 ± 5.117 | 0.4172 | 8.187 ± 4.300 | 94.552 ± 46.844 | 0.4268 | 108.772 ± 56.135 |
| **MCP-1 (CCL2)** | 2.788 ± 3.068 | 0.0248* | 1.332 ± 1.756 | 3.471 ± 3.955 | 0.2136 | 2.428 ± 2.444 | 14.512 ± 9.569 | 0.6911 | 12.777 ± 7.502 |
| Mean ± SD | Statistical tests were two sample T test for parametric and Wilcoxon test for nonparametric data. | | | | | | | | |
| n = 19 for male LyzMyd88$^{-/-}$ N= 16 for Myd88$^{flox/flox}$ control | | | | | | | | | |
| n = 16 for female LyzMyd88$^{-/-}$ and Myd88$^{flox/flox}$ control | | | | | | | | | |
| units: pg/ml/mg dry soleus | | | | | | | | | |
| All *P* values reached Benjamini Hochberg FDR < 0.20 | | | | | | | | | |
| Red denotes signficant effects of Myd88 KD in myeloid cells: * <0.05, **<0.01, ***<0.001 | | | | | | | | | |

**Table S2. Concentrations of cytokines in the muscle baths at the three points of sample collection in the LPS stimulation experiments.** In general, in this experiment, baseline values of cytokines at T0 were extremely low. When comparing the absolute concentrations of muscle cytokines within the baths at each time point, females showed a reduction in IP-10 and KC secretion in the LyzMyd88 KO muscles at $T_2$ and decreases in IL-6 and KC at $T_1$. They also showed reductions in MCP-1 and IP-10 at $T_2$. In contrast, males exhibited reductions in TNF$\alpha$, IL-6, IP10, and MCP1 at $T_0$. At $T_2$, TNF$\alpha$ was marginally decreased and G-CSF marginally increased in the LYZ-Myd88$^{-/-}$ muscles. Muscles from LyzMyd88$^{-/-}$ mice produced lower baseline cytokines (particularly in males) in the KO mice suggesting that some of the baseline cytokine measurements observed may have arisen from the muscle tissue during the brief < 3 min of buffer exposure, and that these were influenced by LyzMyd88$^{-/-}$. None of these single timepoint measurements affected the data reported in the main text which were based on the rate of secretion within intervals.
